# Targeted deletion of the RNA-binding protein *Caprin1* leads to progressive hearing loss and impairs recovery from noise exposure in mice

**DOI:** 10.1101/2021.05.20.444945

**Authors:** Lisa S. Nolan, Jing Chen, Ana Cláudia Gonçalves, Anwen Bullen, Karen P. Steel, Sally J. Dawson, Jonathan E. Gale

**Affiliations:** UCL Ear Institute, 332 Gray’s Inn Road, London. WC1X 8EE; Wolfson Centre for Age-Related Diseases, King’s College London, Guy’s Campus, London. SE1 1UL

**Keywords:** Stress granules, *Caprin1*, hearing loss, neurodegeneration, cochlea

## Abstract

Cell cycle associated protein 1 (Caprin1) is an RNA-binding protein that can regulate the cellular post-transcriptional response to stress. It is a component of both stress granules and neuronal RNA granules and is implicated in neurodegenerative disease, synaptic plasticity and long-term memory formation. Our previous work suggested that Caprin1 also plays a role in the response of the cochlea to stress. Here, targeted inner ear-deletion of *Caprin1* in mice leads to an early onset, progressive hearing loss. Auditory brainstem responses from *Caprin1*-deficient mice show reduced thresholds, with a significant reduction in wave-I amplitudes compared to wildtype. Whilst hair cell structure and numbers were normal, the inner hair cell-spiral ganglion neuron (IHC-SGN) synapse revealed abnormally large post-synaptic GluA2 receptor puncta, a defect consistent with the observed wave-I reduction. Unlike wildtype mice, mild-noise-induced hearing threshold shifts in *Caprin1*-deficient mice did not recover. Oxidative stress triggered TIA-1/HuR-positive stress granule formation in *ex-vivo* cochlear explants from *Caprin1-*deficient mice, showing that stress granules could still be induced. Taken together, these findings suggest that Caprin1 plays a key role in maintenance of auditory function, where it regulates the normal status of the IHC-SGN synapse.

## Introduction

Cell cycle associated protein 1 (Caprin1; also known as RNG105) is a RNA-binding protein that was originally identified as a promoter of cell proliferation (Grill et al., 2004; B. Wang, David, & Schrader, 2005). Subsequently, it has been shown to be a core nucleating component of stress granules (N. Kedersha et al., 2016; Solomon et al., 2007) and also to regulate local RNA translation in neuronal RNA granules (Nakayama et al., 2017; Shiina, Shinkura, & Tokunaga, 2005; Shiina, Yamaguchi, & Tokunaga, 2010). Stress granules are concentrated aggregates of RNA binding proteins and RNA which form by liquid-liquid phase separation under certain types of stress (Alberti, Gladfelter, & Mittag, 2019; N. Kedersha et al., 2016; Riggs, Kedersha, Ivanov, & Anderson, 2020). Caprin1 plays a key role in promoting this condensation mechanism by competing with USP10 for binding to G3BP1 (N. Kedersha et al., 2016, 2020). In a highly dynamic mechanism stress granules recruit specific RNAs, inhibiting translation of these RNAs and either store or degrade them during cellular stress via interactions with processing bodies. Although in recent years many components of stress granules have been identified and the mechanism regulating their formation has been elucidated their precise function is still unclear. It is suggested that stress granules are a pro-survival mechanism, regulating translation of mRNAs and coordinating cell signaling during stress thereby contributing to cellular homeostasis (Buchan & Parker, 2009; Mahboubi & Stochaj, 2017).

In contrast to their suggested role in protecting cells during cellular stress, stress granules have been linked with various pathologies including cancer, viral infections and neurodegeneration (Cao, Jin, & Liu, 2020; L. Chen & Liu, 2017; Mahboubi & Stochaj, 2017). In cancer, assembly of stress granules is associated with resistance to chemotherapy and metastasis (Somasekharan et al., 2015). During viral infection many different viruses have been shown to use various strategies to inhibit stress granule formation designed to avoid stress granule-mediated stalling of viral RNA translation (Riggs et al., 2020). Mutation of stress granule components including TDP-43, TIA1 and FUS is associated with neurodegenerative diseases such as amyotrophic lateral sclerosis and core stress granule components are localized to pathological aggregates in Alzheimer’s disease and Huntington’s disease (Cao et al., 2020; L. Chen & Liu, 2017; Mahboubi & Stochaj, 2017). Accumulation of such aberrant or persistent stress granules in pathological aggregates associated with neurodegeneration has been attributed to the dysregulation of stress granule formation/disassembly and has been implicated in age-related disease (Cao et al., 2020).

Previously, we have identified a role for Caprin1 and stress granules in the response to cochlear stress (Goncalves et al., 2019; Towers, Kelly, Sud, Gale, & Dawson, 2011b). The process of hearing sound is a very sensitive and sophisticated process that itself generates various types of intrinsic cellular stress: mechanical, ionic, oxidative, and synaptic (Poirrier, Pincemail et al. 2010, Someya and Prolla 2010). The cochlea is also subject throughout its life to various kinds of additional extrinsic stresses most notably as a result of damage from environmental agents such as noise and ototoxic drugs (e.g. aminoglycoside antibiotics and cisplatin) (Ohlemiller 2006, Cannizzaro, Cannizzaro et al. 2014). All of these mechanisms can contribute to hearing loss and their effects are cumulative resulting in a common age-related decline in auditory function (Bowl and Dawson 2019). Caprin1-positive stress granules form in sensory hair cells of the mammalian inner ear in response to aminoglycoside antibiotics and pharmacological manipulation of stress granule formation can protect sensory hair cells from aminoglycoside-induced ototoxicity (Goncalves et al., 2019). These data suggest Caprin1 and stress granules play an important role in auditory protection during cellular stress.

Since, *Caprin1* homozygote knockout mice exhibit neonatal lethality due to respiratory failure we created an inner ear conditional *Caprin1* knockout mouse (*Caprin1* cKO), driven by the *Sox10-Cre* allele (S. M. Muller et al., 2008) in which to study the role of Caprin1 and stress granules in auditory protection during cellular stress. We show that *Caprin1* deficient mice still form stress granules in response to arsenite treatment suggesting Caprin1 is not necessary for stress granule formation. However, mice lacking Caprin1 in the inner ear exhibit an early-onset hearing loss that progresses with age and an inability to recover from the effects of mild-noise exposure unlike wildtype mice. Furthermore, we provide evidence that loss of the *Caprin1* protein in mice leads to a defect in the IHC-SGN synapse structure and function, evidenced by a reduction in auditory brain stem response (ABR) wave-I amplitudes and changes in the morphology of the post-synaptic puncta, observed by labelling the GluA2 subunit of the post-synaptic AMPA-type ionotropic glutamate receptors. Our results have implications for our understanding of the role of RNA granule proteins and their role in determining how the auditory system responds and recovers from damage.

## Results

### Generation of *Caprin1* deficient mice

Mice carrying a germline knockout for the *Caprin1* gene are neonatal lethal (Shiina et al., 2010). Therefore, to understand the role of Caprin1 in the auditory system we undertook a conditional knockout approach using a *Sox10-cre* driver specific to inner ear and craniofacial neural crest-derived tissues (Matsuoka et al., 2005). Mice carrying a conditional allele for *Caprin1* were generated at the Wellcome Sanger Institute, U.K. and crossed with mice carrying the *Sox10-cre* transgene to generate conditional knockout *Caprin1^tm3d/tm3d^* (*Caprin1* cKO) mice (Fig.1A). In contrast to the systemic knockout, *Caprin1^tm3d/tm3d^* mice were viable and survived to adulthood. Quantitative PCR confirmed significant knockdown of the Caprin1 mRNA transcript in the brain of *Caprin1^tm3d /tm3d^* mice compared to control mice (p=2E-06), with heterozygote mice having intermediate levels of expression (Fig. 1B). Caprin1 protein expression was compared in the cochleae of *Caprin1^tm3d/tm3d^* and wildtype mice using immunofluorescence (Fig.1C-H). In wildtype mice at P28, Caprin1 expression appears relatively ubiquitous as described for other tissues (Solomon et al., 2007). However, increased immunoreactivity was observed in inner hair cells (IHC) within the organ of Corti (Fig. 1C, arrowhead), and in the spiral ganglion (SG) (Fig.1C, E and G). Labelling of the Deiter’s cells that cradle the outer hair cells (OHCs) was also apparent (arrows in Fig.1 E). In the cochlea of the *Caprin1* cKO mice, Caprin1 immunoreactivity was much reduced (Fig.1D) and the reduction was particularly notable in the IHC region. A very small number of cells appeared to escape the Cre recombination and continued to express Caprin1 protein (see SG in Fig.1D). The gross anatomy and overall structure of the cochlea was not altered in the cKO mice with three rows of outer hair cells and a single row of inner hair cells present in mutant and wildtype mice.

**Fig. 1.**
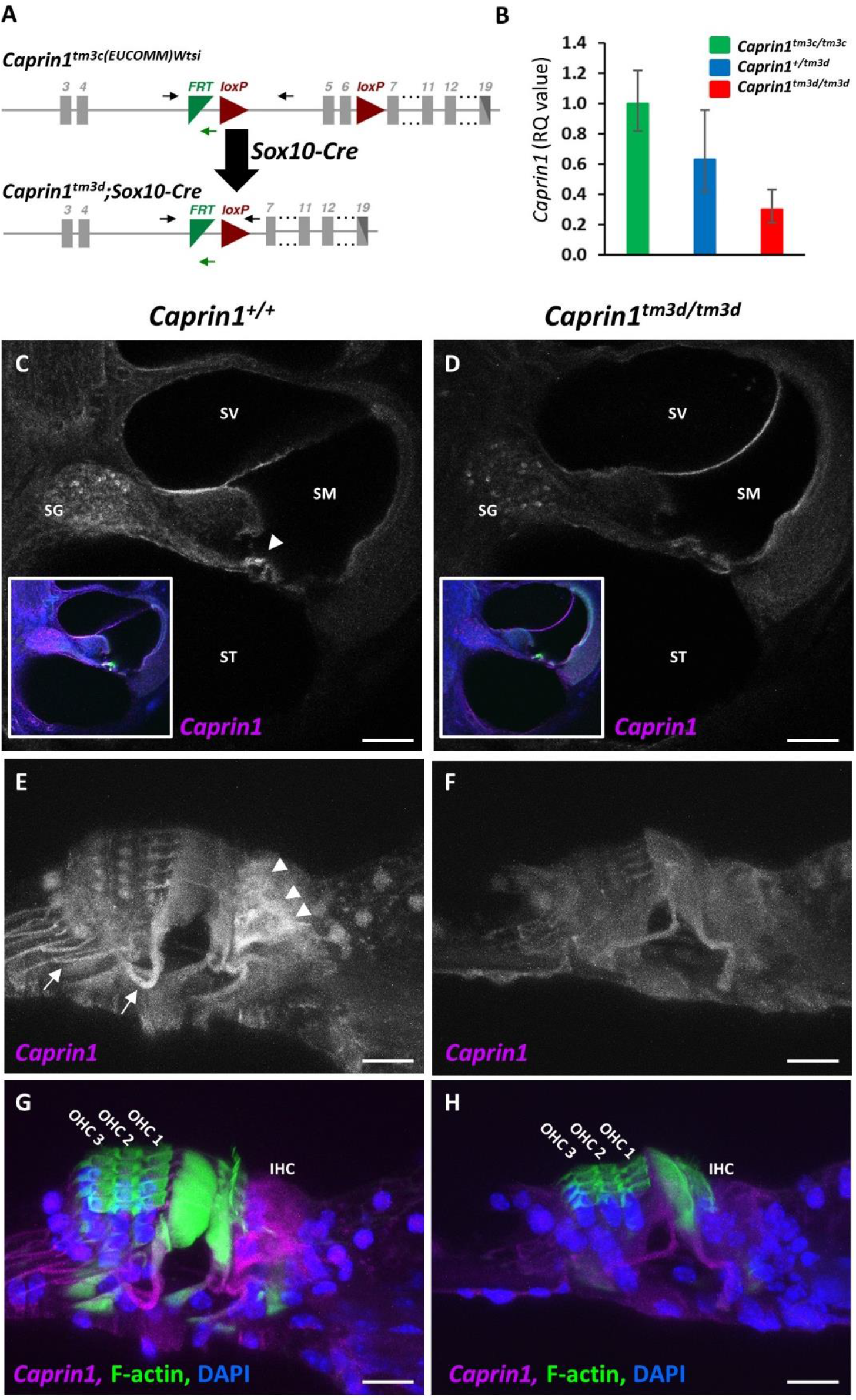
Targeted deletion of *Caprin1* in the inner ear. **A.** Schematic shows the design of the conditional allele for *Caprin1^tm3c(EUCOMM)Wtsi^*. Exons are shown in grey. *LoxP* sites flank the critical exons (exons 5-6) of the *Caprin1* gene. The position of primers *Caprin1_173389_F*, *Caprin1_173389_F* and *CAS_R1_Term* are indicated by black and green arrows respectively. Mice carrying the conditional allele for *Caprin1* were crossed with mice expressing *Cre-*recombinase driven by the *Sox10* promoter deleting the floxed critical exons and generating a frameshift mutation in the inner ear *(Caprin1^tm3d^)*. **B.** Quantitative PCR of *Caprin1* expression in the brain confirmed knockdown of *Caprin1* in *Caprin1^tm3d/tm3d^* mice compared to control mice (p=2E-06, T-test comparing ΔCT values). Heterozygotes displayed *Caprin1* levels intermediate to *Caprin1^tm3d/tm3d^* mice and *Caprin^tm3c/tm3c^* controls. *RQ*, relative quantification adjusted to endogenous control values, error bars show 95% confidence intervals. **C-H.** Immunofluorescence labelling with anti-Caprin1 on vibratome sections of inner ears from P28 wild-type **(C, E & G)** and mutant **(D, F & H)** mice: anti-Caprin1 (magenta; for clarity shown alone in greyscale in C-F); DAPI (blue) and phalloidin staining of F-actin (green). Images are maximum projections of confocal Z-stacks. **C-D.** Low magnification view of the basal cochlear coil, scale bar: 100μm. SV, scala vestibuli; SM, scala media; ST, scala tympani; SG, spiral ganglion. Insert shows the merged image. **C.** In the wild-type cochlea Caprin1 immunoreactivity is predominantly localized to the inner hair cells (arrowhead) and SG. **E-H.** High magnification of the organ of Corti from the mid-cochlear coil, scale bar 10μm. **G, H** show the merge of **E, F**, respectively. **E**. Caprin1 immunoreactivity in the wild-type organ of Corti is detected particularly in IHCs (arrowheads) and also appears concentrated in the Deiter’s cell region (arrows). **F.** Minimal Caprin1 immunoreactivity was detected in the mutant organ of Corti. However, a small number of SG neurons (SGNs) continue to express Caprin1 in the mutant cochlea **(D).**

### Caprin1-deficient mice exhibit early-onset progressive hearing loss

To assess the effect of Caprin1-deletion on hearing function auditory brainstem response (ABR) recordings were performed at successive time-points in a longitudinal hearing study out to postnatal day (P)210 (Fig.2A). At P28, the earliest time-point assessed, hearing thresholds in *Caprin1^tm3d/tm3d^* mice were already significantly elevated across most frequencies compared to wildtype controls (Fig.2A). With increasing age loss of auditory sensitivity in *Caprin1^tm3d/tm3d^* mice progressed significantly in the mid to high-frequencies (18-24kHz), such that by P210 little recordable hearing could be detected at frequencies above 18kHz (Fig.2A). In comparison, a slower loss of auditory sensitivity was observed in the low to mid-frequency range (Fig.2A e.g. 12kHz). Hearing thresholds in heterozygous mice (*Caprin1^+/tm3d^*) did not significantly differ from wildtype at P28 (Fig. 2A). However, at older ages heterozygotes displayed a small, but significant elevation in the high frequencies compared to wildtype mice (Fig.2A).

**Fig. 2.**
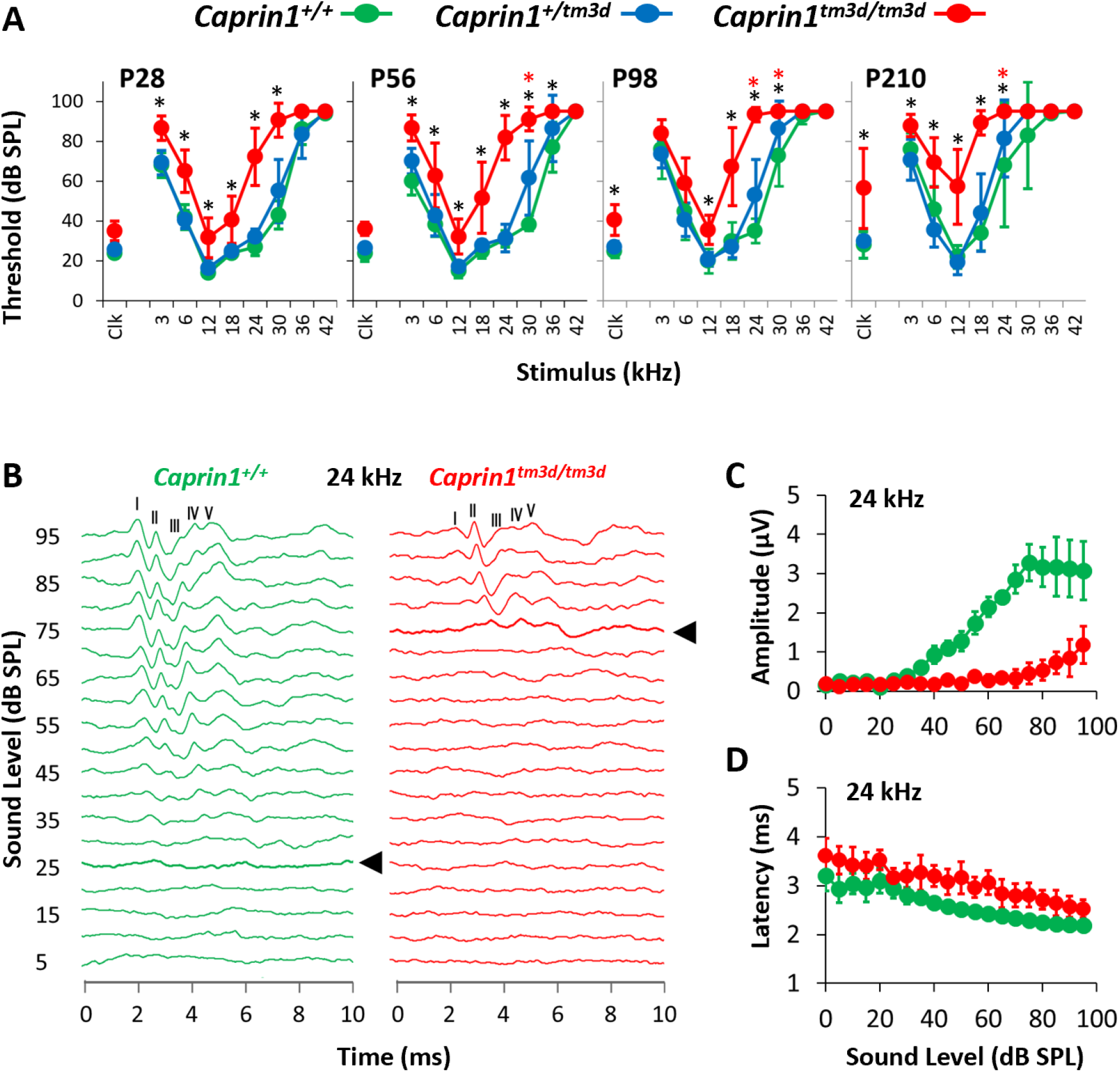
*Caprin1-* deficient mice exhibit a progressive hearing loss. **(A)** Mean ABR thresholds (± SD) for click and pure-tone stimuli for *Caprin1^+/+^* (n=5), *Caprin1^+/tm3d^* (n=7) and *Caprin1^tm3d/tm3d^* (n = 9) mice at P28, P56, P98 and P210 (*Caprin1^tm3d/tm3d^*, n=7 at P210; 2 mice died before P210). * mean hearing thresholds that significantly differ between *Caprin1^+/+^* and *Caprin1^tm3d/tm3d^* mice; * mean hearing thresholds that significantly differ between *Caprin1^+/+^* and *Caprin1^+/tm3d^* mice (p <0.05). **(B)** Representative ABR waveform traces obtained from *Caprin1^+/+^* and *Caprin1^tm3d/tm3d^* mice at P28 in response to a 24kHz pure tone presented from 0 to 95 dB SPL in 5dB increments. The five positive peaks of the ABR waveform trace are numbered I to V. The ABR threshold is denoted by the thick line and arrowhead: 25dB SPL versus 75dB SPL in *Caprin1^+/+^* and *Caprin1^tm3d/tm3d^* mice, respectively. **(C)** The growth of ABR wave I amplitude and **(D)** the change in ABR wave I latency as a function of the sound pressure level (dB SPL) in *Caprin1^+/+^* (n=5) and *Caprin1^tm3d/tm3d^* (n = 6) mice at P28 in response to 24kHz stimuli is shown. **(C)** Amplitudes (μV) and **(D)** latencies (ms) are plotted as mean values; error bars: ± SD.

A more detailed analysis of ABR waveforms at 24kHz (Fig. 2B) the frequency at which *Caprin1^tm3d/tm3d^* mice exhibited a large significant elevation in hearing thresholds at P28 (27 ± 4 vs 70 ± 16 dB SPL, mean ± SD, *Caprin^+/+^* vs *Caprin1^tm3d /tm3d^* respectively), showed that the amplitudes of ABR waves I and III-V were reduced in *Caprin1^tm3d/tm3d^* mice. We quantified this reduced amplitude focusing on ABR wave I which in humans and most mammals is generally agreed to reflect the action potentials in auditory nerve fibres driven by activity at the IHC-SGN synapse (Jewett, Romano, & Williston, 1970; Parkkonen, Fujiki, & Makela, 2009; Willott, 2006). In *Caprin1^tm3d/tm3d^* mice, ABR wave I amplitudes were greatly reduced compared to wildtype mice and this reduction became more pronounced as a function of increasing sound level (Fig.2C). In parallel, a small increase in the latency of ABR wave I was observed in *Caprin1^tm3d /tm3d^* mice compared to wildtype (Fig.2D).

### *Caprin1*-deficient mice exhibit a post-synaptic defect

To characterise the functional deficit that underlies the hearing loss in *Caprin1*-deficient mice we used both cellular and sub-cellular markers in whole mount preparations of the organ of Corti. The cochlea is organized in a tonotopic gradient with low frequencies detected towards the apex and higher frequencies towards the base. We focused our investigations on the mid-basal cochlear coil at P28, which includes the 24kHz region, a region which exhibited a severe functional deficit at this time-point (Fig. 2A). First, we examined the general architecture of the organ of Corti. Phalloidin staining of filamentous actin (F-actin) revealed that the epithelial surface appeared normal with no obvious abnormality in the gross morphology of the organ of Corti in *Caprin1^tm3d/tm3d^* mice compared to wildtype mice (Fig.3A, B). Organisation of both IHC and outer hair cell (OHC) rows appeared normal (Fig.3A, B) and labelling with an antibody to Myosin7a revealed normal appearance of IHC cell bodies (Fig.3E, F). Similarly, scanning electron microscopy showed normal appearance of stereociliary hair bundles in *Caprin1^tm3d/tm3d^* mice (Fig.3Q). Quantification of IHCs and OHCs revealed no difference in the number of sensory hair cells between *Caprin1^tm3d/tm3d^* and wild-type mice (Fig.3O). We then examined the IHC synapse using antibodies to CtBP2 to label pre-synaptic IHC ribbons (Khimich et al., 2005), and to GluA2 to label post-synaptic AMPA-type ionotropic glutamate receptors on adjacent afferent dendrites of type I spiral ganglion neurons (SGNs) (Matsubara, Laake, Davanger, Usami, & Ottersen, 1996). The number of CtBP2-positive puncta and GluA2-positive puncta per IHC were reduced in *Caprin1^tm3d /tm3d^* mice compared to wildtype; this difference was not statistically significant but was at the border of statistical significance for CtBP2 (p = 0.05 for CtBP2 and p = 0.12 for GluA2). We then compared the mean cross-sectional area of the CtBP2- and GluA2-positive puncta and found that the GluA2-positive post-synaptic densities were almost 2-fold larger in *Caprin1^tm3d/tm3d^* mice compared to wildtypes (Fig.3N: 0.47 SD± 0.03μm^2^ versus 0.24 SD± 0.03μm^2^, respectively; p = 0.0001). We did not observe a difference in size of the pre-synaptic CtBP2-positive puncta between *Caprin1^tm3d /tm3d^* and wild-type mice (Fig.3M). However, immunofluorescence images suggested that there was a consistent decrease in the expression of nuclear localized CtBP2 in IHC of *Caprin1^tm3d/tm3d^* compared to wildtype mice (Fig.3A-D, G-H). Quantitative analysis of the relative intensity of the CtBP2 fluorescent signal in IHC nuclei and synaptic ribbons compared to the IHC cytoplasm in *Caprin1^tm3d/tm3d^* and wildtype mice showed the CtBP2 fluorescent signal was significantly reduced in IHC nuclei of *Caprin1^tm3d/tm3d^* mice whereas the relative level of the expression in the synaptic ribbons was similar between wildtype and *Caprin1* cKO mice (Fig.3P, Fig.S1).

**Fig. 3.**
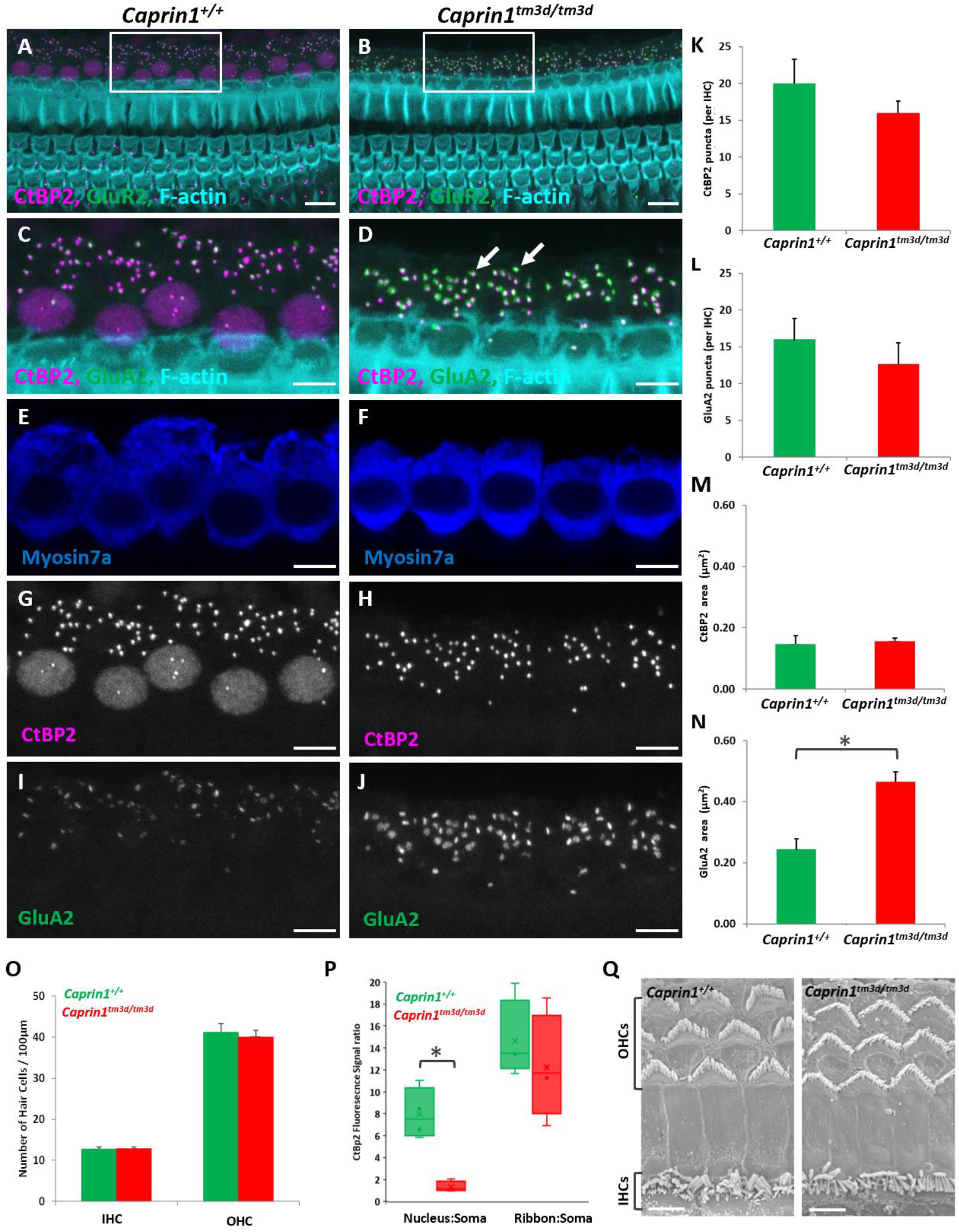
Caprin1-deficient mice exhibit a post-synaptic defect. (**A-B**) Maximum projection images of immunofluorescence confocal Z-stacks from the mid-basal cochlear coil (24kHz region at P28) reveal the normal F-actin pattern (F-actin rich hair bundles, cuticular plates and pillar cell surfaces) of the reticular lamina in *Caprin1^+/+^* and *Caprin1^tm3d/tm3d^* mice. The boxed regions in **A** and **B** are shown in (**C,E,G,I**) and (**D,F,H,J**). Labelling with anti-CtBP2 and anti-GluA2 reveals alterations in the inner hair cell synapses with significantly enlarged GluA2 puncta (arrows) in *Caprin1^tm3d/tm3d^* (**C-D**). There are no obvious abnormalities in the IHCs, labelled with anti-Myosin7a (**E,F**). Quantification of images from the mid-basal cochlear coil (24kHz region) at P28 in *Caprin1^+/+^* (n = 4) and *Caprin1^tm3d/tm3d^* (n = 5) mice reveals no significant difference between the number of pre-synaptic CtBP2-positive ribbons (**K**) or the number of post-synaptic GluA2 receptor puncta (**L**) per IHC. The cross-sectional area of IHC pre-synaptic ribbons was not affected (**M**) however there was a significant difference in the size of post-synaptic GluA2 receptors between *Caprin1^+/+^* and *Caprin1^tm3d/tm3d^* mice, * p= 0.0001. (**O**) The number of IHCs and OHCs per 100 μm length of the organ of Corti was the same in *Caprin1^+/+^* and *Caprin1^tm3d/tm3d^* mice. (**P**) *Caprin1^+/+^* mice showed a typical intensity profile of the CtBP2 immunofluorescence signal across in the IHC soma (see **G** and Suppl Fig. 1) from minimal expression in the cytoplasm (soma) increasing to *8× greater signal in the nucleus and *14× greater signal in the ribbon puncta (i.e. the ribbon puncta expression is *1.75× greater than the nuclear expression). The nuclear CtBP2 expression was much reduced in *Caprin1^tm3d/tm3d^* IHCs (significant difference from *Caprin1^+/+^* *p=0.009) whereas the average intensity of the ribbon puncta was not significantly different from *Caprin1^+/+^.* (**Q**) SEM micrographs of the surface of the organ of Corti reveals normal morphology in *Caprin1^tm3d/tm3d^* i.e with v-shaped stereociliary hair bundles in OHCs and the ‘balustrade’-like hair bundles of IHCs. Scale bar: 2μm, images taken from the mid-basal cochlear coil. Error bars in all graphs represent 1xSD. Scale bars in **A-B**, 10μm; **C-J** 5μm and **Q**, 2μm.

### *Caprin1* deficient mice do not recover from noise-induced hearing loss

Given the progressive nature of the hearing loss in the *Caprin1^tm3d/tm3d^* mice and Caprin1’s role in regulating translation during cellular stress, we tested whether these mice are more vulnerable to stress, in the form of acoustic trauma. We exposed *Caprin1^tm3d/tm3d^* mice to two different noise exposure paradigms (Fig.4). The first is designed to induce a temporary threshold shift (TTS) (Hesse et al., 2016; Kujawa & Liberman, 2009) from which wildtype mice are known to recover (Fig.4A, C). The second paradigm is designed to induce a permanent threshold shift (PTS) (Amanipour et al., 2018; Y. Wang, Hirose, & Liberman, 2002) from which wildtype mice would not completely recover (Fig.4B).

**Fig. 4.**
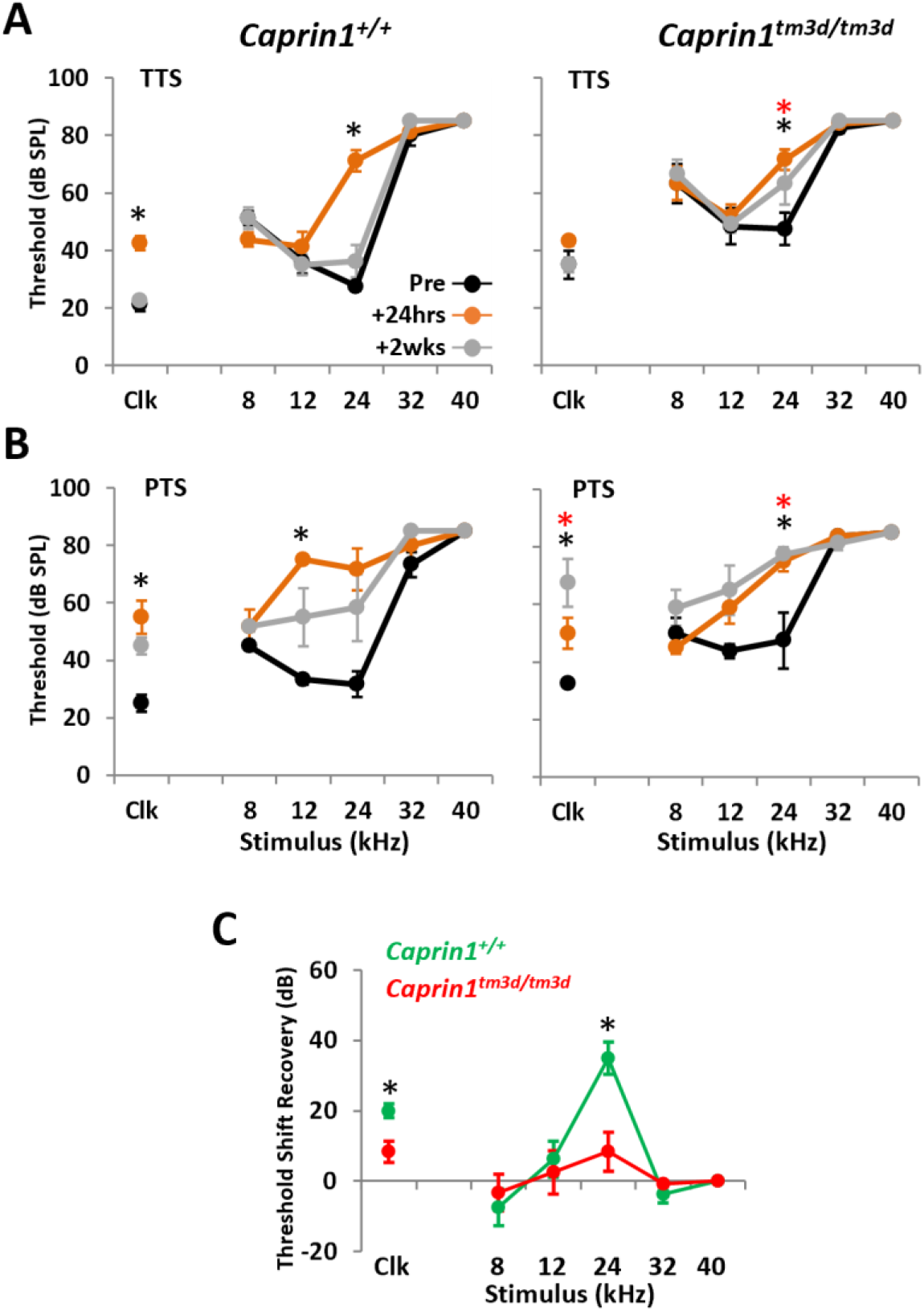
*Caprin1* deficient mice do not recover from noise exposure. **A&B.** ABR thresholds for broadband click (Clk) and tone pip stimuli (8-40kHz) for *Caprin1^+/+^* (n=4 in A, n=3 in B), and *Caprin1^tm3d/tm3d^* (n = 6 in A, n = 4 in B) mice following a temporary threshold shift (TTS, 8-16kHz octave-band noise, 100 dB SPL, 2 hours) ***in A***, or permanent threshold shift (PTS, 8-16kHz octave-band noise, 110 dB SPL, 3 hours) noise exposure regime ***in B***. ABRs were measured pre-noise exposure and at 24hrs and 2wks following noise exposure. * and * denote thresholds that are significantly elevated (p<0.05) at 24hrs and 2 weeks post-noise exposure, respectively. **C.** Recovery of ABR threshold shifts for *Caprin1^+/+^* and *Caprin1^tm3d/tm3d^* mice following TTS regime (calculated by subtracting the threshold at 24hrs from that at 2wks). *\** denotes threshold shifts that significantly differ (p<0.05) between *Caprin1^+/+^* and *Caprin1^tm3d/tm3d^* mice. All data are mean ± SEM.

ABR thresholds in wildtype and *Caprin1^tm3d/tm3d^* mice were significantly elevated 24hrs following the TTS paradigm, particularly at 24kHz (Fig.4A). However, by 2 weeks post-exposure the elevated ABR thresholds in wildtype mice had recovered (Fig.4A). In contrast, auditory sensitivity had not fully recovered in *Caprin1^tm3d/tm3d^* mice after the same 2-week period. The mean ABR thresholds at 24kHz remained significantly elevated at 2 weeks compared to pre-exposure measures; 63 ± 7 vs 48 ± 34 dB SPL, respectively; p=0.0421 (Fig.4A). Subtraction to reveal the threshold shifts confirmed recovery of hearing thresholds in wildtype mice, but not *Caprin1^tm3d/tm3d^* mice (Fig.4C).

The TTS paradigm induces a TTS that predominantly localizes to the 24kHz region of the cochlea (Fig.4D). To investigate the response of *Caprin1^tm3d/tm3d^* cochlea to greater stress we increased the extent of the acoustic trauma to a PTS paradigm (Fig.4B). The PTS paradigm induces a substantial threshold shift in the mid-frequency region of the cochlea. ABR thresholds in wildtype and *Caprin1^tm3d/tm3d^* mice were elevated across 12-24kHz and in response to broadband click stimuli 24 hours following exposure (Fig.4B, see Clk). However, two weeks later ABR thresholds in wildtype mice were still elevated although they had recovered such that they did not significantly differ from pre-exposure levels (Fig.4B). In comparison, the elevated ABR thresholds in *Caprin1^tm3d/tm3d^* mice show little evidence of recovery and remained significantly elevated, even appearing to worsen after 2 weeks (Fig.4B, see Clk). These results suggest that the absence of Caprin1 protein in the cochlea impairs the ability of the cochlea to recover from acoustic stress.

To further understand why *Caprin1^tm3d/tm3d^* mice are unable to recover from noise exposure, we compared the effect of noise exposure on Caprin1 protein expression and localisation in the wildtype and cKO cochlea. Two weeks following the TTS noise exposure paradigm pronounced Caprin1 immunoreactivity was detected in both the IHC region (Fig.5A, C, thick arrow) and the SGNs (Fig.5E, G) in the wildtype cochlea. In noise-exposed cochleae, we observed clusters of Caprin1 immunoreactivity localized to cytoplasm above the nucleus of the OHCs (thin arrow, Fig.5A, C) and discrete punctate regions of Caprin1 immunoreactivity were detected in the organ of Corti consistent with the appearance of stress granules (Fig.5A, C short arrow). Negligible Caprin1 immunoreactivity was observed in the organ of Corti (Fig.5B, D) or SGN (Fig.5F, H) of *Caprin1^tm3d/tm3d^* mice after noise exposure consistent with its inner ear cKO status. However, as described previously (Fig.1C), a small fraction of SGN cells in noise-exposed *Caprin1^tm3d/tm3d^* mice continue to display Caprin1 immunoreactivity suggesting these cells escape Cre-targeted deletion (Fig.5F, H, arrows).

**Fig. 5.**
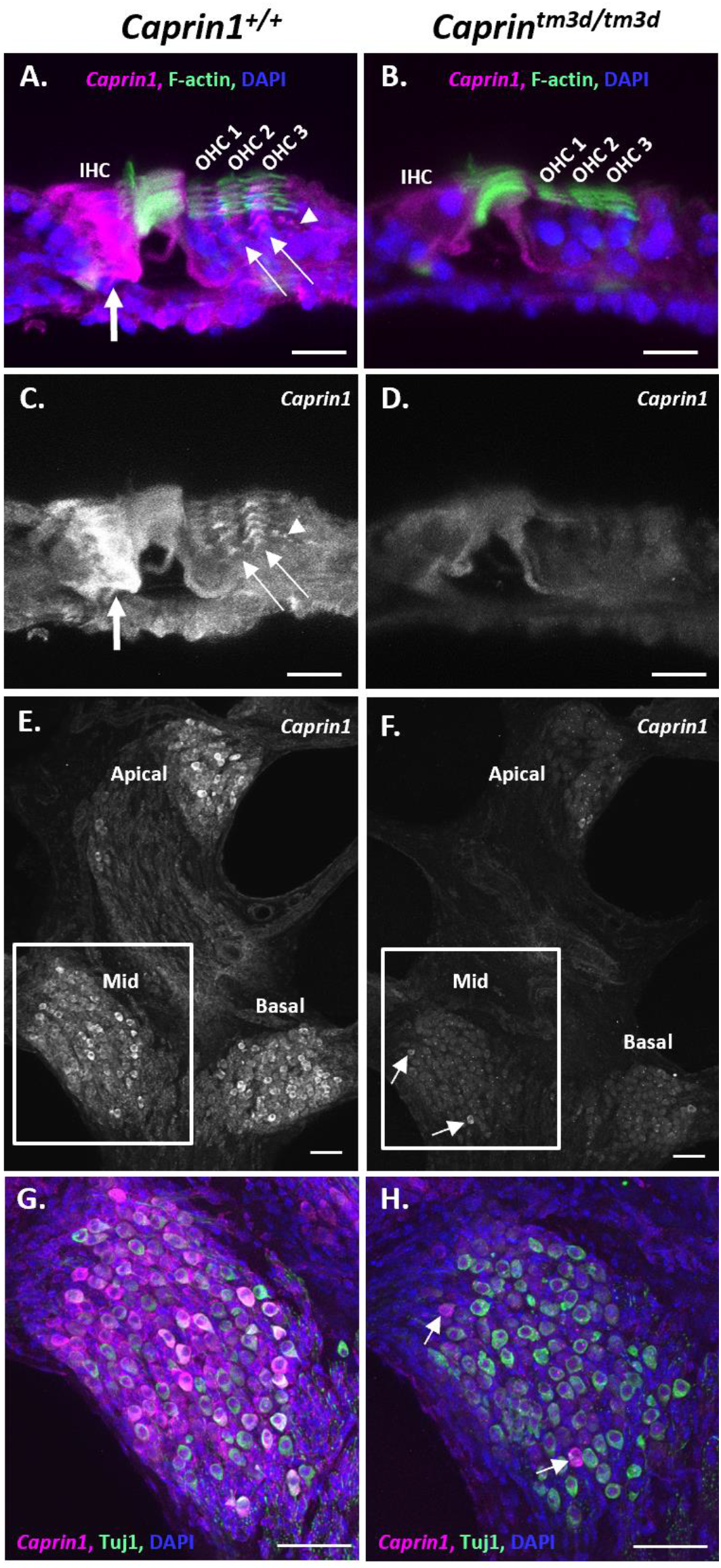
Expression of Caprin1 in the cochlea after noise exposure. **A-H.** Results of immunofluorescence labelling with anti-Caprin1 on vibratome sections of wild-type (**A, C, E & G**) and cKO (**B, D, F & H**) mouse organ of Corti (**A-D**) and SGN (**E-H**) at P44 two weeks following a TTS noise exposure regime. All images are representative confocal maximum projections. Anti-Caprin1 (white in **C-F**; magenta in the merge **A, B, G, H**); anti-Tuj1 labelling predominantly SGNs (green, **G-H**), DAPI (blue) and phalloidin labelling of f-actin (green, **A-B**). **A-D.** The organ of Corti in the apical cochlear coil. **A, B.** Show the merged image of **C, D,** respectively. In the wild-type cochlea (**A, C**) abundant Caprin1 immunoreactivity is localized to the IHC region (thick arrow) and is also concentrated above the OHC nuclei (thin arrow, **A, C**). Evidence of punctate regions of Caprin1 immunoreactivity were also detected (arrowhead, **A, C**). Minimal Caprin1 immunoreactivity was detected in the organ of Corti of *Caprin1^tm3d/tm3d^* mice (**B, D**)**. E-F,** a mid-modiolar vibratome section shows Caprin1 labelling of SGNs. The boxed regions are shown in **G-H,** respectively. In the wild-type cochlea (**E & G**) Caprin1 immunoreactivity is localized to the cytoplasm of the majority of SGNs, across all cochlear coils, with some cells expressing more than others. In the *Caprin1^tm3d/tm3d^* cochlea, Caprin1 is not observed in SGNs (**F & H**), except for a few neurons that continue to exhibit Caprin1 immunoreactivity (**F & H**, white arrows). IHC: inner hair cell; OHC: outer hair cell. Images represent data from at least 3 mice. Scale bars: 20 μm in **A-F**, 50 μm in **G-H**.

### *Caprin1*-deficient mice can still form stress granules

We have shown that *Caprin1^tm3d /tm3d^* mice fail to show normal recovery from two different acoustic stress paradigms when measured two weeks following noise exposure. This may be due to the significantly altered IHC-SGN post-synaptic morphology identified in *Caprin1^tm3d/tm3d^* mice but it could also be due to an effect of Caprin1 knockdown on stress granule formation or function. The appearance of discrete aggregates of Caprin1-immunoreactivity in the organ of Corti of noise-exposed wild-type mice (Fig.5) suggests that stress granule formation may play a role in auditory protection during noise exposure. Therefore, we tested the ability of the *Caprin1^tm3d /tm3d^* cochlea to form stress granules.

Previously we have shown that Caprin1-containing stress granules are induced in response to various types of cellular stress (Goncalves et al., 2019; Towers et al., 2011b). Aminoglycoside treatment triggers the formation of Caprin1-containing stress granules in *ex-vivo* postnatal rat cochlear explants (Towers et al., 2011b) and *in-vivo* in mouse hair cells (Goncalves et al., 2019). Sodium arsenite, used to mimic oxidative stress, induces the formation of Caprin1-containing stress granules in UB/OC-2 cells (a mouse inner ear sensory epithelial cell line) and in *ex-vivo* cochlear explants (Goncalves et al., 2019). Caprin1-containing stress granules colocalize with polyA+ mRNA and known stress granule markers such as TIA1 cytotoxic granule-associated RNA-binding protein (TIA-1) whereas in some cases Caprin1-positive granules do not contain TIA-1 (Goncalves et al., 2019; Towers et al., 2011b). Although Caprin1 has been shown to be a key regulator of stress granule formation and its over-expression is sufficient to induce stress granule formation, it is unclear whether it is an absolute requirement (N. Kedersha et al., 2016; Towers, Kelly, Sud, Gale, & Dawson, 2011a). Hence, we investigated whether stress granules could still form in *Caprin1^tm3d/tm3d^-*cochlea in response to cellular stress using two markers of stress granules, TIA-1 and Human antigen R (HuR) (N. L. Kedersha, Gupta, Li, Miller, & Anderson, 1999; Riggs et al., 2020).

Sodium arsenite was applied to *ex-vivo* postnatal mouse cochlear explants to compare stress granule formation in *Caprin1^tm3d/tm3d^* and control (*Caprin^tm3c/tm3c^*, *sox10-ve*) mice (Fig.6). Cochlear explants from control mice exhibited robust stress granule formation in response to 1 hour of sodium arsenite (Fig.6). Abundant TIA-1-positive and Caprin1-positive (Fig.6A) stress granules were observed in the OHC region and somewhat less robustly in the IHC region of control mice. HuR-positive stress granules followed a similar distribution pattern but appeared less widespread (Fig.6B) than the Caprin-positive stress granules. Many TIA-1 and Caprin1-positive stress granules were observed (Fig.6A, arrows) and a similar pattern was observed for Caprin1 and HuR-positive granules (Fig.6B, arrows). In comparison, in the *Caprin^tm3d/tm3d^* cochlea although a similar distribution pattern of TIA-1-positive stress granules was observed compared to that of control mice (Fig.6A, B), HuR-positive stress granules appeared reduced (Fig.6C, D). Moreover, in the *Caprin^tm3d/tm3d^* cochlea the TIA-1-(Fig.6A, arrowheads) or HuR-(Fig.6B, arrowheads) positive stress granules did not contain Caprin1, consistent with the conditional knockout phenotype of the *Caprin^tm3d/tm3d^* mice. These data also suggest that stress granule formation is not dependent on Caprin1.

**Fig. 6.**
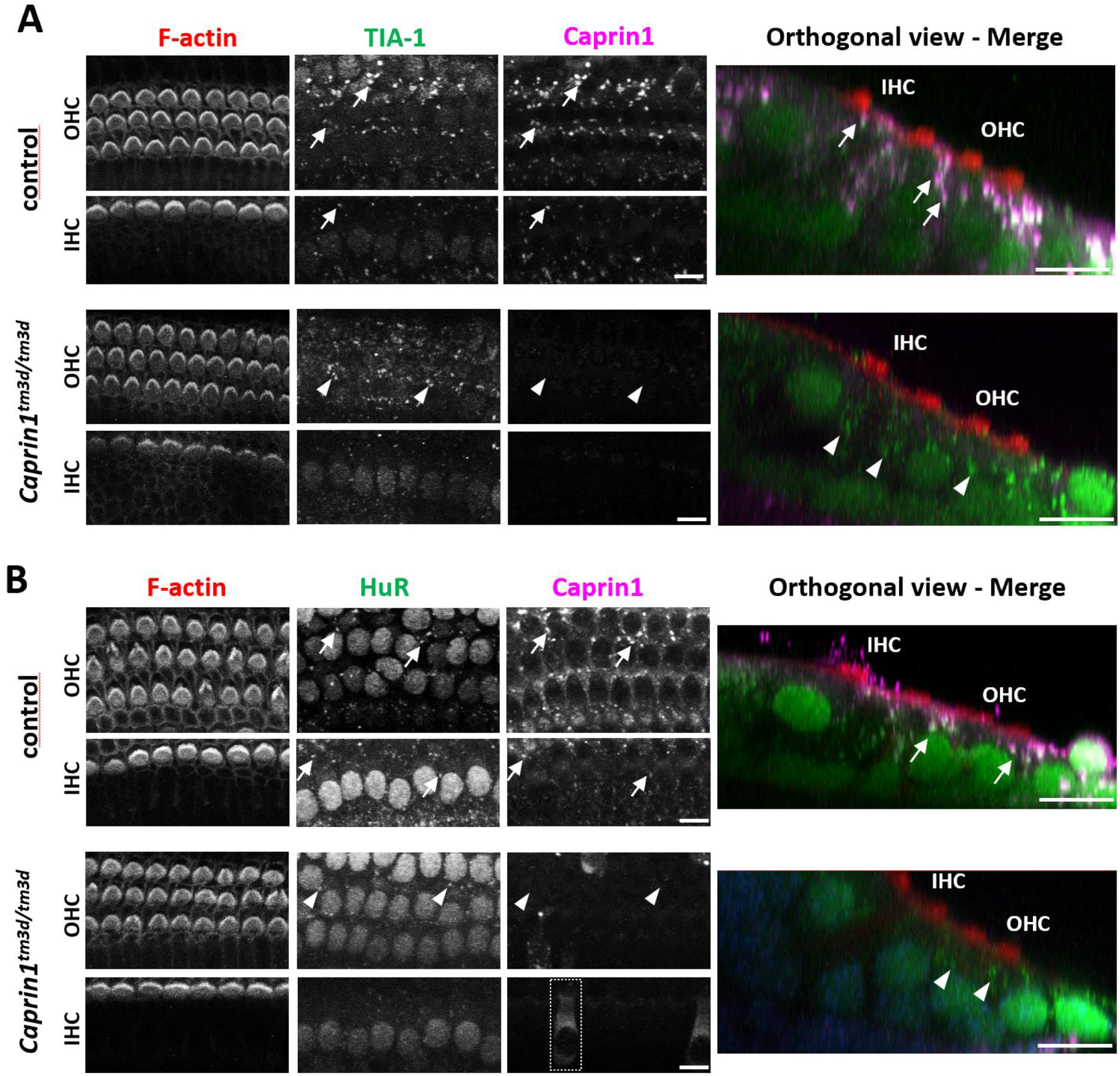
*Caprin1*-deficient mice can still form stress granules. *Ex-vivo* postnatal cochlear explants from *control* (see methods) and *Caprin1^tm3d/tm3d^* mice all treated with 0.5mM sodium arsenite for 1 hr to induce stress granule formation. (**A**) Immunofluorescence maximum projections of confocal image stacks showing expression of Caprin1 and TIA-1 (to label stress granules). *Caprin1^tm3d/tm3d^* mice show minimal expression of Caprin1 in the organ of Corti. In both control (arrows) and *Caprin1^tm3d/tm3d^* (arrowheads) explants TIA-1-positive stress granules form in the cells (including IHCs and OHCs) in response to arsenite treatment. In controls, Caprin1 and TIA-1 colocalise in the majority of stress granules. (**B**) Maximum projections of confocal image stacks showing expression of *Caprin1* and an alternative stress granule marker HuR. The same pattern is observed with arsenite treatment, with HuR-positive/Caprin1-positive stress granules (arrows in controls) and HuR-positive stress granules (arrowheads) still formed in the *Caprin1^tm3d/tm3d^* hair cells (and other cell types). However, as in the *Caprin1^tm3d/tm3d^* SGN (see Fig.1C), a few OHCs and IHCs in the *Caprin^tm3d/tm3d^* cochlea continue to exhibit Caprin1-immunoreactivity consistent with Caprin1 protein expression again suggesting a few cells escape the *Sox10-Cre* recombination (boxed region shows an example of an IHC). All explants (minimum n = 3 per experiment) were from the mid-basal cochlear coil and images were acquired from the mid-region of the explant. Scale bar, 10 μm for all.

## Discussion

### The stress granule RNA-binding protein Caprin1 is essential for maintenance of hearing

Stress granules are thought to play a protective role during cellular stress and dysregulation of their formation and disassembly has been implicated in a number of pathological processes, most notably neurodegeneration (Cao et al., 2020; L. Chen & Liu, 2017; Mahboubi & Stochaj, 2017). However, to date, our knowledge of stress granule biology is largely based on experiments performed in yeast or from manipulation of stress granule protein expression in various cell lines, using heat shock or sodium arsenite to induce stress granule formation. There is little corroboration of these findings from *in-vivo* work, especially in mammals in relation to native physiological stress. A limitation to these investigations has been that generation of knockout mutant mice for stress granule components such as Caprin1, G3BP1 and TIA-1 leads to embryonic or neonatal lethality (Martin et al., 2013; Piecyk et al., 2000; Shiina et al., 2010). Therefore, generation of the conditional knockout *Caprin1^tm3d/tm3d^* mouse model here represents one of the first opportunities to investigate the role of a stress granule protein in a mature mammalian system that is susceptible to physiological stress, the inner ear. Here, we report that conditional knockout of the stress granule RNA-binding protein Caprin1 in the inner ear leads to an early onset and progressive hearing loss in mice suggesting that it plays an important role in the protection and maintenance of the cochlea. The simplest explanation for the hearing loss we observed in *Caprin1^tm3d/tm3d^* mice is that the abnormal post-synaptic morphology we describe manifests itself as a defect in synaptic signaling between the IHC and the SGNs that is measured as a reduction in the ABR wave I and consequent overall hearing thresholds.

In the absence of Caprin1, we find that cells within the cochlea can still generate stress granules in response to oxidative stress *in vitro* and to noise exposure *in vivo* but we find that they are less able to recover from auditory stress in the form of noise exposure. To our knowledge this is the first report to demonstrate that conditional deletion of an RNA-binding protein and stress granule component leads to an increased susceptibility to extrinsic, cellular stress *in vivo*.

Under normal conditions, in the absence of an applied extrinsic stress, Caprin1 is widely expressed in cochlear cells in the adult mouse inner ear consistent with the ubiquitous expression reported in other tissues (Solomon et al., 2007). However, some cells show a more intense staining suggesting increased Caprin1 expression in the IHC region and SGNs (Fig. 1c). In the presence of extrinsic stress including noise and oxidative stress we found punctate staining of Caprin1 in hair cells and in SGNs in response to noise in wildtype mice (Figs. 5 and 6). The punctate staining co-localizes with stress granule markers TIA-1 and HuR suggesting stress granule formation occurs in these cells in response to both types of stress. This is consistent with data showing that Caprin1-positive granules form in response to ototoxic aminoglycoside antibiotic exposure (Gonçalves et al., 2019; Towers et al., 2011a). At the outset, we hypothesised that Caprin1 and thus potentially stress granules, play an important homeostatic, protective role in maintenance of sensory receptor hair cells during different types of cellular stress. In support of this, in the absence of Caprin1 within the inner ear, *Caprin1^tm3d/tm3d^* mice have a high-frequency hearing loss at 4 weeks that is progressive, and extends into lower-frequencies with increasing age (Fig. 2A), a pattern of hearing loss which is commonly found in patients with age-related hearing loss. Heterozygous *Caprin1^+/tm3d^* mice display similar hearing thresholds to *Caprin1^+/+^* mice except for a small elevation of thresholds at higher frequencies, although this does not progress markedly with age (Fig 2A). Characterization of the cochlea of *Caprin1^tm3d/tm3d^* mice at P28 when they already have a significant hearing loss found no obvious loss of either IHC or OHC and the cellular architecture of the organ of Corti appeared normal. These data indicate that the hearing loss does not result from a gross developmental defect and is more consistent with a deficit in auditory maintenance or protection.

### Evidence that the IHC-SGN synapse is altered in *Caprin1^tm3d/tm3d^* mice

Reduced ABR wave I amplitudes in the presence of an otherwise normal audiogram is a characteristic feature of cochlear synaptopathy; a neuropathy whereby the synaptic connections between IHCs and the peripheral afferent dendrites of the SGNs are lost or damaged in response to noise exposure and/or the ageing process (Kujawa & Liberman, 2009, 2015; Liberman, 2017; Wu et al., 2019). ABR wave I amplitudes were reduced in *Caprin1^tm3d/tm3d^* mice from P28, the earliest time-point assessed (Fig. 2) consistent with a cochlear synaptopathy. After detailed investigation using pre and post synaptic markers, we found that the GluA2 post-synaptic densities were abnormal, with GluA2-positive cross-sectional areas that were nearly twice as large in *Caprin1^tm3d/tm3d^* mice compared to wildtype (p < 0.0001). We did not find a significant difference in the number of GluA2-labelled post-synaptic densities in IHCs between *Caprin1^tm3d/tm3d^* mice and wildtype mice (Fig. 3). The values are consistent with the numbers reported in the literature for normal hearing mice; approximately 17 synapses per IHC for the mid-cochlear coil (Kujawa & Liberman, 2009). Additionally, there was no significant difference in the number of CtBP2-labelled pre-synaptic ribbons in IHCs, although we note that the reduction was at the borderline of significance (p = 0.05) suggesting further investigation of the pre-synaptic ribbons is warranted, as this may be related to the GluA2 morphology difference. However, unlike the post-synaptic elements, the average size of the CtBP2-labelled pre-synaptic ribbons was unchanged in cKO mice compared to wildtype.

GluA2 is a crucial component of the AMPA-type ionotropic glutamate receptor, the major excitatory synaptic receptor in the auditory system, crucial for transmission of sound-induced activity from the cochlea to the auditory cortex (Reijntjes & Pyott, 2016; Takago & Oshima-Takago, 2018). As well as being a key regulator of stress granule formation Caprin1 has been implicated in regulation of local RNA translation in neuronal RNA granules located at synaptic junctions (Sephton & Yu, 2015; Shiina et al., 2005; Shiina et al., 2010). In those studies, loss of Caprin1 in hippocampal neurons led to loss of synaptic strength and a consequential deficit in long term memory formation in mutant mice. Using genome wide profiling of mRNA localization, Shiina, Yamaguchi et al. 2010, assigned the cause of this deficit to a lack of localization of dendritic mRNAs to synapses in the absence of Caprin1, resulting in changes in the distribution of synaptic proteins, particularly AMPA receptors, consistent with the changes we observe here. Combining those data with our own data from *Caprin* cKO mice described here suggests that Caprin1 is responsible for regulating the local translation of critical synaptic proteins and hence maintaining the integrity and function of the primary auditory synapse. We suggest that when Caprin1 expression is knocked down or absent a critical homeostatic control mechanism becomes dysregulated leading to changes in post-synaptic structure and function, the latter being consistent with the observed reduction in wave 1 amplitudes in cKO mice (Fig.2c). In support of this homeostatic requirement, a unilateral reduction in sound-induced activity in the immature mouse cochlea (at p11) has previously been shown to modulate the size and the molecular make-up of the IHC-SGN synapse (Barclay, Constable, James, Thorne, & Montgomery, 2016). Similar to our observations in *Caprin1^tm3d/tm3d^* mice, the effect was restricted to the changes in the size of the post-synaptic densities labelled with GluA antibodies (Barclay et al., 2016).

A further striking and consistent difference that we observed in cKO mice was the loss of CtBP2 staining in the IHC nuclei of *Caprin1^tm3d/tm3d^* mice. The *CtBP2* gene is subject to alternative splicing producing both short (*CtBP2-S*) and long (*CtBP2-L*) isoforms, in addition to a third isoform termed Ribeye (Schmitz, Konigstorfer, & Sudhof, 2000; Stankiewicz, Gray, Winter, & Linseman, 2014). Ribeye localizes to ribbon synapses present in sensory cell-types and is critical to their function, including the IHC synapse (Schmitz et al., 2000; Stankiewicz et al., 2014). In contrast, *CtBP2-L* contains a nuclear localization signal, and both *CtBP2-S* and *CtBP2-L* function as corepressors by interacting with transcription factors to repress their activity (Stankiewicz et al., 2014). Currently, aside from Ribeye, little is known regarding the expression and function of the additional *CtBP2* isoforms in the cochlea. Our observations that *Caprin1^tm3d/tm3d^* mice exhibit a loss of nuclear CtBP2 expression suggests the presence of a nuclear isoform of the *CtBP2* gene in the cochlea that is modulated, directly or indirectly by Caprin1. How this loss of nuclear CtBP2 expression manifests itself at the level of the IHC and thus contributes to the auditory defect in *Caprin1^tm3d/tm3d^* mice remains to be determined.

### Stress granule formation in *Caprin1^tm3d/tm3d^* mice and its role in auditory protection

Environmental stress in the form of day-to-day sounds, and at the more extreme, noise exposure, is arguably the biggest extrinsic stressor to the inner ear and a major contributor to age-related hearing loss (Keithley, 2019; Liberman, 2017). Intense noise exposure over hours damages hair cell stereocilia and leads to hair cell death and a permanent hearing loss (Y. Wang et al., 2002). However, moderate noise exposure typically results in a reversible hearing loss, with recovery of ABR thresholds *but* a persistent cochlear synaptopathy and reduced wave-I, which has been termed ‘hidden hearing loss’, since noise-exposed (or aged) humans may have normal pure-tone audiogram thresholds but have significant difficulties distinguishing speech sounds in background noise (Kujawa & Liberman, 2009, 2015; Schaette & McAlpine, 2011). Here, we found that unlike wildtype mice, ABR thresholds in *Caprin1* cKO mice fail to recover from such moderate noise exposure. Using that same noise paradigm, we observed enhanced expression of Caprin1 in the IHC region in wildtype mice, but not in the cKO. These data taken together suggest that Caprin1 may be necessary for auditory protection during stress and/or for repair and recovery from stress-induced damage. Our data indicate that the failure of cKO mice to recover from noise exposure does not result from an inability to form stress granules since cKO cells were still able to do this (Fig.6). Whether the absence of Caprin1 might alter the efficacy of the overall stress granule response remains to be determined but, since different RNA binding proteins are known to bind and recruit different populations of specific mRNAs to RNA granules and stress granules, it seems likely that the absence of Caprin1 would alter the mRNA components of stress granules (Khong et al., 2017; Tian, Curnutte, & Trcek, 2020). Our present data suggest that Caprin1 plays a key role within stress granules or neuronal RNA granules and is a key determinant of the cochlea’s homeostatic response to noise exposure. The data also suggest that the regulation of cochlear synaptic form and function is dependent on neuronal RNA granules and the function of Caprin1. Additionally, Caprin1 may regulate other molecular functions of IHCs, potentially through its effects on nuclear CtBP2. In future work it will be important to characterize the nature of the mRNA molecules recruited to cochlear RNA granules by Caprin1 during stress. In summary, the data suggest a key role for Caprin1 acting pre- or post-synaptically to determine how the auditory system responds to and recovers from damage, revealing a potential therapeutic avenue for preventing acquired hearing loss.

## Materials and Methods

All experiments involving animals in this study were performed in accordance with the regulations of the U.K. Animals (Scientific Procedures) Act of 1986 (ASPA). Experiments were carried out under UK Home Office licences, and the study was approved by the Ethical Review Committees for: King’s College London, the Wellcome Sanger Institute and University College London. Mice were maintained in individually-ventilated cages, subject to a twelve-hour light/dark cycle under standard humidity and temperature conditions in a specific pathogen-free environment. Mice were culled using methods approved under these licences to minimize any possibility of suffering.

### Generation and genotyping of Caprin1 conditional knockout mice

Mice carrying a conditional-ready allele for *Caprin1* (*Caprin1^tm3c(EUCOMM)Wtsi^*; MGI ref: 5692641) on a C57BL/6N; C57BL/6N-*A^tm1Brd/a^* genetic background were generated at the Wellcome Sanger Institute by the Mouse Genetics Project (Skarnes et al., 2011; White et al., 2013). Flp recombinase-mediated excision of the cassette inserted into the *Caprin1^tm3a^* allele was used to generate the *Caprin1^tm3c^* allele which has LoxP sites flanking exons 5-6 of the *Caprin1* gene, upstream of the known functional domains of the *Caprin1* gene (Fig. 1A; https://www.mousephenotype.org/data/genes/MGI:1858234).

In order to delete the floxed exons of *Caprin1* in the inner ear to produce *tm3d* mice, *Caprin1^tm3c^* mice were mated to *Sox10-Cre* mice (*Tg(Sox10-cre)1Wdr;* MGI ref: 3586900; a gift from Prof. William Richardson, UCL, U.K.). The *Sox10* gene is expressed early in development around embryonic day 9.5 in the otic vesicle before specification of the pro-sensory domain and in neural crest-derived cells (Kuhlbrodt, Herbarth et al. 1998, Pusch, Hustert et al. 1998, (J. Chen et al., 2014; S. M. Muller et al., 2008). Cre-*lox*P recombination generates the *Caprin1^tm3d^* allele, deleting exons 5 and 6 in the inner ear (Matsuoka, Ahlberg et al. 2005). The *Sox10-Cre* mice used to breed with the *Caprin1^tm3c^* mice were maintained on a mixed genetic background including C57BL/6N and CBA. Aberrant homologous recombination has been reported when the *Sox10-Cre* transgene is transmitted through the paternal germline (Crispino, Di Pasquale et al. 2011). Therefore, experimental *tm3d* animals were generated by crossing female *Caprin1^tm3d/+^* or *Caprin1^tm3d/tm3d^* mice with *Sox10-Cre* negative males *Caprin1^tm3c/+^* or *Caprin1^tm3c/tm3c^*. *Caprin1^tm3d/tm3d^* mice were initially bred and maintained in the laboratory of Prof. Karen Steel (Wolfson Centre for Age-Related Diseases, King’s College London, U.K.) and used for ABR recordings in a longitudinal hearing study (Fig.2). Subsequent experiments were performed with *Caprin1^tm3d/tm3d^* mice bred and maintained in the laboratories of Dr Sally Dawson and Prof. Jonathan Gale (Ear Institute, University College London, U.K.). Experimental controls used were either littermates that were wildtype for the *Caprin1^tm3c^* allele in the presence of the *Sox10-Cre* transgene (*Caprin1*^+/+^ mice) or did not carry the *Sox10-Cre* allele (*Caprin1^tm3c/tm3c^* mice). Both male and female animals were used in experiments. Mice were born in normal Mendelian ratios and no excessive deaths were recorded in any genotype group. Details of numbers of experimental animals used are in the figure legends.

Genotyping was carried out by PCR using DNA extracted from pinna tissue. PCRs were run in singleplex using primer pairs: *Caprin1_173389_F* 5’-AGCCAGTGCTCTTTGAACCC-3’ and *Caprin1_173389_R* 5’-GCCAAACATCCACCACTGAC-3’ which generate a 600bp product at the native genomic locus and a 752bp product in the presence of floxed allele for *Caprin1*; *Caprin1_173389_F* and *CAS_R1_Term* 5’-TCGTGGTATCGTTATGCGCC-3’ which generate a 236bp product in the presence of the floxed allele for *Caprin1* (see Fig. 1); and *Sox10Cre_F* 5’-GCGGTCTGGCAGTAAAAACTATC-3’ and *Sox10Cre_R* 5’-GTGAAACAGCATTGCTGTCACTT-3’ which generate a 101bp product in the presence of the *Sox10-Cre* transgene. Mice carrying the *Caprin1* deletion (*Caprin1^tm3d^* allele) were confirmed by presence of the 236bp PCR band (*Caprin1^tm3c^* allele) in conjunction with the 101bp band (*Sox10-Cre* allele). PCR cycling conditions are available upon request.

### Quantitative real-time PCR

Brain tissue from *Caprin1^tm3d /tm3d^*, *Caprin1^+ /tm3d^* and *Caprin1^tm3c /tm3c^* (Sox10-negative) mice was dissected in RNAlater (QIAGEN^®^) (n=3 individual animals per genotype), lysed and homogenized with a TissueRuptor and total RNA was isolated with a RNeasy^®^ Plus Mini Kit (QIAGEN^®^). cDNA synthesis was performed using the Omniscript^®^ Reverse Transcription Kit (QIAGEN^®^) with random primers (Promega) and rRNasin^®^ Plus Ribonuclease Inhibitor (Promega). Parallel cDNA synthesis reactions were run for each sample minus the Omniscript Reverse Transcriptase as a negative control for gDNA contamination. *Caprin1* expression was determined by qPCR using a Taqman gene expression assay from Applied Biosystems (Rn01512768-m1) on a SDS7500 Fast PCR System (Applied Biosystems). Relative quantification of *Caprin1* was based on triplicate samples using the 2^−*ΔΔ*Ct^ method with eukaryotic 18S rRNA as the endogenous control (Applied Biosystems, Cat. No: 4319413E).

### Preparation of cochleae and immunofluorescence analysis

Auditory bullae were dissected and fixed in 4% paraformaldehyde (PFA) for 1 hour at room temperature, washed in phosphate-buffered saline solution (PBS) and decalcified in 4.13% EDTA, pH7.4, in PBS for 72 hours at 4°C. For vibratome sections, bullae were mounted in 4% low melting point agarose (Sigma-Aldrich, Gillingham, UK) and sectioned at 200 μm using a 1000 Plus Vibratome (Intracel, Royston, UK). For preparation of cochlear whole mounts, the organ of Corti was dissected into 4 half-coils (apex; mid-apical; mid-basal and basal) using an adaptation of the Eaton-Peabody Laboratories protocol. In brief, the organ of Corti was bisected along the modiolus following the half-coil of the apex aligned to the oval window as a guide. The lateral wall and stria vascularis were carefully removed from each half coil as described (Eaton-Peabody Laboratories, 2017). Vibratome slices were permeabilized and blocked with 0.5% Triton-X 100, 10% goat serum in PHEM buffer (60mM PIPES, 25mM Hepes, 10mM EGTA, 2mM MgCl_2_, pH 6.9) for 2 hours at room temperature. Organ of Corti whole mounts were permeabilised with 5% tween-20 in PBS for 1 hour at room temperature and blocked with 10% horse serum, 0.5% Triton-X 100 in PBS for 2 hours at room temperature. Primary antibody incubations were performed at 4 °C overnight. After PBS washes, secondary antibody incubations were either performed for 1 hour (whole mounts) or 2 hours (vibratome sections) at room temperature in the dark. F-actin, abundant in hair cell stereocilia, was labelled with 10nM phalloidin-Atto 647N (Sigma-Aldrich, Gillingham, UK) and nuclei were visualized with 1 μM DAPI. Imaging was performed with a Zeiss LSM 510 or 880 confocal microscope using 10× (0.45 N.A.), 20× (0.8 N.A.), and 63× oil (1.4 N.A.) objectives and Zen 3.0 SR software (Black edition). Saturation of images and zero black levels were avoided in image capture. For display, any adjustments were made equally to images from *Caprin1^tm3d/tm3d^* and wildtype mice. In taking the confocal settings used to image Caprin1 in unexposed wildtype mice, we found that for the noise-exposed cochlea there was significantly enhanced labelling of Caprin1 such that we needed to reduce laser excitation (% laser power) by half, keeping all other settings the same, indicating an overall increase in Caprin1 expression. *Ex-vivo* cochlear culture explants were immunostained as described previously (Goncalves et al., 2019). Cochlear explants were imaged with a Zeiss 510 NLO multi-photon upright confocal system using a 63× (1.0NA) immersion objective. All immunostaining experiments used a minimum of 3 mice per genotype and representative images are shown in the figures. All images are maximum intensity projections of confocal z-stacks.

### Antibodies

#### Vibratome sections

Primary antibodies: rabbit anti-Caprin1, 1:200 (Proteintech Europe, #15112-1-AP); mouse (IgG_2a_) anti-TuJ1, 1:500 (Covance, #MMS-435P). Secondary antibodies: goat anti-(rabbit IgG) conjugated to Alexa Fluor 633, 1:500 (#A-21070); goat anti-(mouse IgG_2a_) conjugated to Alexa Fluor 488, 1:500 (#A-21131) – both Invitrogen. *Organ of Corti whole mounts.* Primary antibodies: mouse (IgG1) anti-CtBP2, 1:400 (BD Biosciences, #612044); mouse (IgG_2a_) anti-GluA2, 1:200 (Millipore, #MAB397); rabbit anti-Myosin7a, 1:200 (Proteus Biosciences, #25-6790). Secondary antibodies: goat anti-(mouse IgG1) conjugated to Alexa Fluor 568, 1:500 (#A-21124); goat anti-(mouse IgG_2a_) conjugated to Alexa Fluor 488, 1:500 (#A-21131); goat anti-(rabbit IgG) conjugated to Alexa Fluor 405, 1:300 (#A-31556) – all Invitrogen. *Ex-vivo cochlear cultures.* Primary antibodies: rabbit anti-Caprin1, 1:500 (Proteintech Europe, #15112-1-AP); goat anti-TIA-1 (C-20), 1:300 (SantaCruz Biotechnology, #sc-1751); mouse (IgG1) anti-HuR (3A2), 1:500 (SantaCruz Biotechnology, #sc-5261). Secondary antibodies: donkey anti-(rabbit IgG) conjugated to Alexa Fluor 647, 1:100 (#A-31573, Thermofisher); donkey anti-(goat IgG) conjugated to Alexa Fluor 488, 1:1000 (#A-11055, Thermofisher).

### Hair cell counts and IHC ribbon synapse analysis

Maximum intensity projections were made of confocal Z-stacks collected at 0.5μm intervals (63× oil, 1.4 N.A. objective). The start Z position for the image stack was determined by the first synaptic marker visible in the IHC region (ie at the lowest position) and the end Z position was the top-most OHC stereocilia bundle. Data were collected from the 24kHz region in the mid-basal cochlear coil according to the mouse tonotopic frequency-place map (M. Muller, von Hunerbein, Hoidis, & Smolders, 2005). For the hair cell counts, data were quantified in a 112μm lengths of the organ of Corti (see Fig. 3a for representative section) and Myosin7a and phalloidin were used to identify and count hair cells using the Cell Counter plugin (NIH, Image J software version 1.52d). Data are presented as the number of IHCs or OHCs per 100μm length of the organ of Corti.

For analysis of the IHC ribbon synapse, a region of interest (ROI) measuring 40.3 × 21.3μm was positioned over the IHC region to capture five individual IHCs. Subsequent analysis of the IHC ribbon synapse was performed with Image J software version 1.52d. Each ROI file was imported as greyscale using the Bioformats Importer plugin to split each ROI into four individual channels encompassing: CtBP2; GluA2; Myosin7a and phalloidin immunostaining. To count the number of CtBP2-positive puncta per IHC, the threshold tool with the MaxEntropy pre-set function was used to select all CtBP2-positive puncta and generate a binary image. Counts and cross-sectional area measurements were made using the Analyze Particles function in ImageJ. The total number of CtBP2-positive puncta was then divided by the number of IHCs. Counts and cross-sectional area measurements for GluA2-positive puncta were obtained in the same manner. The use of the maximum projection image for quantification was validated by cross-checking data quantified on a slice-by-slice basis from two wildtype and two *Caprin1^tm3d /tm3d^* samples.

### Scanning electron microscopy (SEM)

Preparation of the organ of Corti for SEM has been described previously (Bullen et al., 2019). In brief, cochleae were fixed in 2.5% glutaraldehyde in 0.1M cacodylate buffer with 3 mM CaCl_2_ for 2 hours at room temperature and decalcified for 48 hours in 4% EDTA at 4°C. The organ of Corti was dissected from decalcified cochlear tissue, post-fixed in OsO4 and processed through the thiocarbohydrazide-Os-repeated procedure (Davies & Forge, 1987). Processed samples were dehydrated in a graded ethanol series, critical point-dried and sputter coated with platinum. Samples were examined in a JEOL 6700F SEM operating at 5kV by secondary electron detection. Imaging was carried out using SEM Supporter software (System In Frontier, Japan).

### Auditory brainstem response

Repeated ABR recordings were measured in *Caprin1^+/+^*, *Caprin1^+/tm3d^* and *Caprin1^tm3d/tm3d^* mice at P28 (P27-29), P56 (P56-57), P98 (P98-101) and P210 (P210-213). Two *Caprin1^tm3d/tm3d^* mice died before the final ABR measurement at P210. Mice were anaesthetised with a cocktail of ketamine (100 mg/kg, i.p., Ketaset) and xylazine (10 mg/kg, i.p., Rompun) and subcutaneous needle electrodes were inserted on the vertex (active) and over the left (reference) and right (ground) bullae. Free-field click (0.01-ms duration) and tone pip stimuli presented at 3, 6, 12, 18, 24, 30, 36 and 42 kHz (5-ms duration, 1-ms rise/fall time) over a range of intensity levels from 0 to 95dB sound pressure level (SPL, 5 dB steps) were performed as described previously and thresholds were defined by the lowest sound intensity giving a visually-detectable ABR response (Ingham et al., 2019). *Caprin1^tm3d/tm3d^* mice and wildtype controls, bred and maintained at the UCL EI, underwent ABR recordings at P28 to confirm the auditory phenotype prior to use of the cochleae tissue in immunofluorescence investigations. Click and tone pip stimuli (8, 12, 24, 32 and 40kHz) were presented from 0-85dB SPL in 5 dB steps using TDT System 3 equipment and software (Tucker-Davis Tech., Alachua, FL) as described previously (Mistry, Nolan, Saeed, Forge, & Taylor, 2014). This ABR protocol was also used for mice that underwent recordings as part of the noise exposure experiments. Mice were recovered between successive ABR recordings with atipamezole (1 mg/kg, i.p. Antisedan). There were no sex-specific differences in hearing thresholds at any time-point for the conditional KO mice (data not shown). Therefore, in all investigations we grouped data from both male and female mice.

Additional input-output analyses were performed using the 24kHz pure-tone ABRs at P28 from the longitudinal hearing study. ABR waveforms were analysed further to determine the amplitude and latency of ABR wave I, using a customised MatLab script. In three *Caprin1^tm3d/tm3d^* mice no wave 1 was detected; these were not included in the analyses.

### Noise exposure

Mice were anesthetized with a cocktail containing ketamine (100mg/kg weight, i.p., Vetalar) and medetomidine (0.83mg/kg weight, i.p., Domitor). A bolus of buprenorphine (0.1mg/kg weight, s.c., Vetergesic) was administered separately for additional analgesia. Saline (0.005ml/kg weight, s.c.) was administered for hydration and eyes protected with Viscotears^®^. Mice at P30 were exposed to 8-16kHz octave-band noise at 100dB SPL for 2 hours to generate a temporary threshold shift (TTS) (Hesse et al., 2016; Kujawa & Liberman, 2009). In a separate experiment, *Caprin1^+/+^* and *Caprin1^tm3d/tm3d^* mice were exposed to 8-16kHz octave-band noise at 110dB SPL for 3 hours to generate a permanent threshold shift (PTS) (Amanipour et al., 2018). Noise exposures was performed in a custom sound-proof booth with an RX6 processor (Tucker-Davis Technologies, TDT) as described previously (Hesse et al., 2016). Pedal-reflex and breathing rate were checked every 30 minutes. As before, mice were recovered at the end of the noise exposure with atipamazole (1mg/kg, s.c.). ABR recordings were performed pre-noise exposure at P28, and then sequentially at 24hours, 1 week and 2 weeks post-exposure. Cochleae were harvested after the final ABR recording for use in subsequent immunofluorescence analysis.

### Cochlear cultures

Cochlear explants from postnatal day 3 pups were dissected and cultured as described previously (Towers et al., 2011b). Experiments were performed on explant from the mid-basal cochlear coil. Cochlear explant cultures were allowed to acclimatize overnight (at least 16 hours) before the addition of 0.5mM sodium arsenite for 1 hour to induce oxidative stress as described previously (Goncalves et al., 2019). Following the sodium arsenite treatment paradigm, cochlear explants were fixed in 4% PFA in PBS for 30 minutes at room temperature, washed 3 times in PBS prior to immunostaining.

### Statistics

Sample sizes were chosen based on our experience with comparable mutant mouse experiments (J. Chen et al., 2014). The unpaired Student t-test was used to compare two independent variables. The paired t-test was used to compare matched samples when ABR recordings were performed on the same animal pre- and post-noise exposure. Data was analyzed with GraphPad Prism 7.0d. For the initial longitudinal hearing study, ABR thresholds were not normally distributed, so the data were first transformed using the arcsine transformation and then analysed using separate linear models for each frequency with a compound symmetric covariance structure and restricted maximum likelihood estimation (Duricki, Soleman, & Moon, 2016). This allowed the inclusion of all available data points, including data from mice that died (2 out of 21) before the final ABR measurement (Krueger & Tian, 2004). For each stimulus the double interaction of genotype and age was measured followed by a Bonferroni correction for multiple testing. The arcsine transformation and mixed model linear pairwise comparison were performed with SPSS v25. For all statistical analyses a p-value <0.05 was considered significant.

## Acknowledgements

We thank Alberto Capurro for writing the Matlab script used in analyzing waveforms and Lorenzo Preite for colony management. This work was supported by grants from the MRC [MR/N004329/1 to SD and JG; G0300212 to KPS] and the Wellcome Trust [091092/Z/09/Z to SD and JG; 100669, 098051 to KPS]. ACG was supported by an Action on Hearing Loss PhD Studentship [EI:595 to JG and SD].

## Author Contributions

*Lisa S. Nolan:* acquisition, analysis, visualization, data interpretation, writing – original draft, writing – review and editing.

*Jing Chen:* acquisition, analysis, visualization.

*Ana-Claudia Goncalves:* acquisition, analysis, visualization.

*Anwen Bullen:* acquisition, analysis, visualization.

*Karen P. Steel:* supervision, funding acquisition, writing – review and editing.

*Sally J. Dawson:* conceptualization, acquisition, analysis, supervision, funding acquisition, visualization, data interpretation, writing – original draft, writing – review and editing.

*Jonathan E. Gale:* conceptualization, acquisition, analysis, supervision, funding acquisition, visualization, data interpretation, writing – original draft, writing – review and editing.

## Conflict of Interest Statement

The authors have no known conflicts of interest to declare.

## Supplementary Figure

**Suppl Figure:**
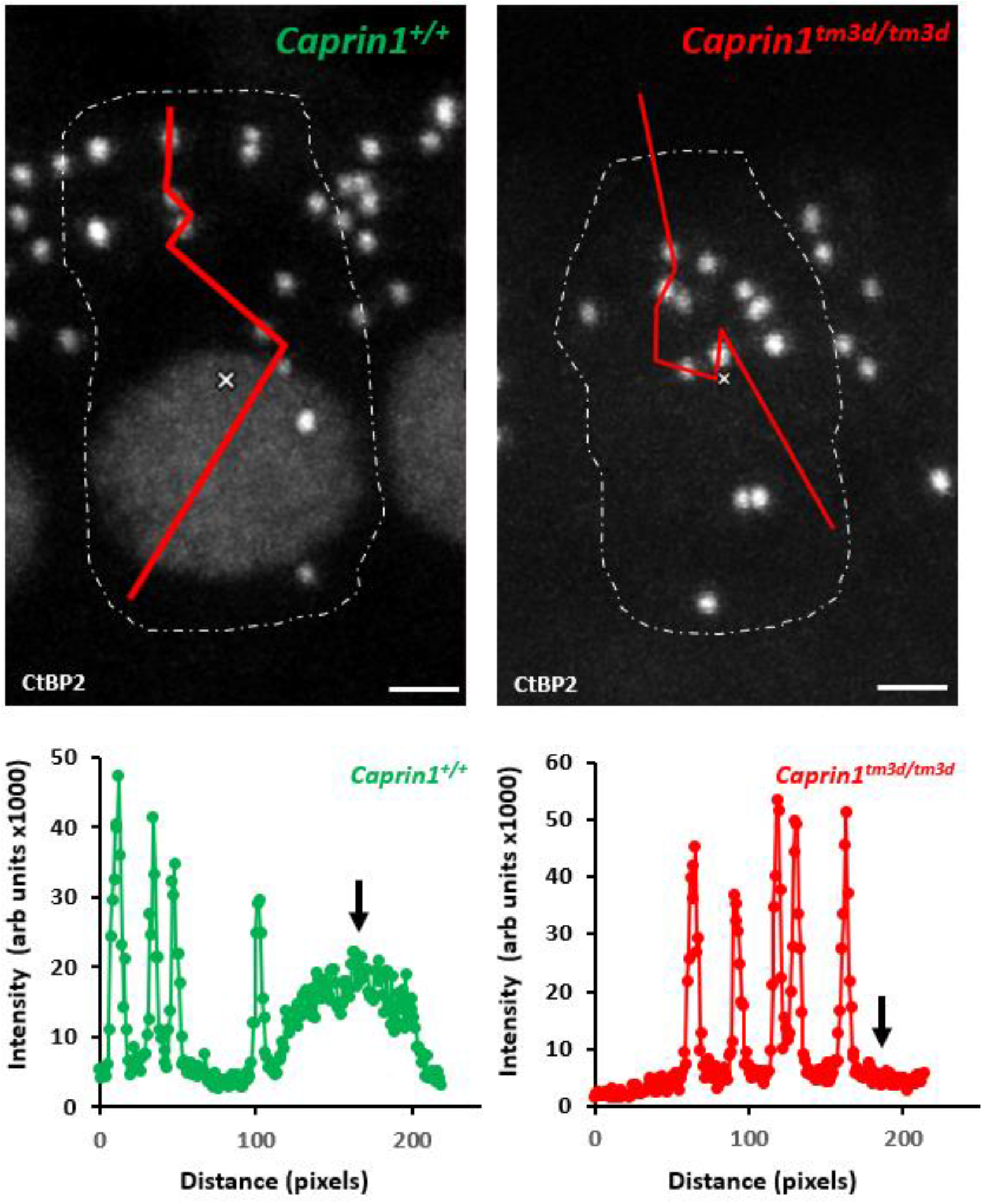
Nuclear CtBP2 expression is reduced in the IHCs of *Caprin1^tm3d/tm3d^* mice. Upper panels, max-projection images of the IHC region from wildtype *Caprin1^+/+^* and *Caprin1^tm3d/tm3d^* mice. The approximate area of a single IHC is outlined based on Myosin 7a expression. The red lines indicate a path drawn through the cell to measure the variation in CtBP2 intensity across the cell (plotted in the graphs in the lower panels). The nuclear region is indicated by the black arrow. Scale bars, 2 μm.

## Notes

### Competing Interest Statement

The authors have declared no competing interest.

